# Are carbohydrate ring distortions determined by biology or crystallography? The impact of experimental conditions on structural data

**DOI:** 10.64898/2026.09.30.755651

**Authors:** Douglas Silva de Lima, Vinícius Ávila Cabral, Hugo Verli

## Abstract

Carbohydrates are essential molecules involved in a wide range of biological processes. While their structural representation in the Protein Data Bank (PDB) has been hindered due to their highly diverse and flexible nature, advancements on X-ray crystallography and CryoEM are accelerating the obtaining of their experimental structures, which in turn supports improved analyses of their structural properties, such as the role of their non-chair conformations on biological processes. In this context, the current work provides a comprehensive and up to date overview of carbohydrate structures within the PDB through classification, quantification, and characterization of monosaccharides and glycan chains, incorporating Cremer-Pople puckering coordinates to evaluate pyranose conformations. Our findings show that, even with a steep increase in CryoEM structures including more complex oligosaccharides, carbohydrate data remains concentrated in small glycan chains, with a about 4% of non-chair conformations in top resolution structures. Free energy landscapes for the pyranose ring in the presence and absence of nine different binding proteins, including both enzymes and non-enzymes, reinforce the preferred stability of chairs and the rarity of conformational induction upon binding.

## Introduction

Carbohydrates are among the most complex and abundant families of biomolecules in nature, playing various roles in biotic systems, such as nutrition,^1^ cell recognition,^2^ immune response,^3^ regulation of the microbiome^4^ and participation in diseases such as diabetes^5^ and cancer.^6^ The simplest and yet highly chemical diverse carbohydrate units are monosaccharides, which can be bonded through variable and flexible glycosidic links to form di-, oligo-, or polysaccharides, which in turn can still be linear or branched. Each glycosidic bond can be oriented in stereoisomeric configurations *α* and *β*, while it can also be frequently associated with lipids and proteins, forming glycolipids and glycoproteins that play essential roles in tissues and organs.^7,8^ Still, in spite of their biological importance, carbohydrates have posed a series of challenges to structural biologists and computational scientists for decades.

The occurrence of errors in protein structures obtained from X-ray crystallography has been recognized for almost 40 years.^9^ Carbohydrates, with their chemical diversity and flexibility, have been recognized in subsequent years for their unique challenges in 3D structure determination and unambiguous annotation in databases.^10,11^ In addition to problems with glycosidic linkages,^12^ later works brought to light geometric distortions of monosaccharides as structural anomalies in the PDB and their potential relation to structure quality and refinement,^13^ despite the probable role of these distorted conformations in enzyme catalysis.^14^ However, there is no indication of the potential biological roles of carbohydrates’ distorted conformations when complexed with noncatalytic proteins.

Unfortunately, extrapolations of these initial analyzes using up-to-date information from the PDB have been severely hindered in the past by a significant number of errors and inconsistencies in the glycan components of these 3D structures, including problems with nomenclature, stereochemical and regiochemical inaccuracies, unlikely conformations, incorrect N-glycan core structures, and poorly modeled glycans, even for high resolution structures.^13,15^ Attempting to circumvent such limitations, our group published in 2020 a global analysis of the PDB, from which a quantitative estimate of the limitations of this database in terms of carbohydrate complexity (that is, the size of polysaccharide chains) and diversity (that is, the types of monosaccharide units), as well as the analysis of puckering residues in the largest dataset so far.^11^ In the following year, a modernized representation of carbohydrate molecules in the PDB was published,^16^ representing a significant leap to support more accurate and systematic analyzes of 3D experimental data on carbohydrates and their relation to biological phenomena.

In this sense, following the continuous growth of entries and improvement in structural quality within PDB, as well as the recent advancement of cryo-electron microscopy (CryoEM) while employing the standardization of carbohydrate representation brought by Shao and co-workers,^16^ the current work aims to deepen previous analyzes of carbohydrate structural content in PDB, searching for evidence for the structural basis for monosaccharide distortions in PDB and its potential impact on multiple biological processes. Accordingly, our results demonstrate that: i) 9% of all carbohydrate structures in the PDB adopt high energy conformations. ii) the quality of the experimental structure remains one of the key determinants for ring distortions in the PDB, an effect similar for enzymes and non-enzymes; iii) CryoEM structures have a propensity up to three times lower to be associated with carbohydrates distorted states, pointing to the role of experimental conditions in residue distortion; iv) such a degree of distortion is also similar among all carbohydrate-binding protein families represented in the database, instead of being a specific phenomenon associated with catalysis; and v) few proteins seem to be able to change the conformational landscape of hexopyranoses towards non-chair states. Taken together, these observations substantially restricting the potential relevance of these unusual conformations to biological phenomena.

## Results and discussion

### PDB Overview

In this study, all mmCIF files from the PDB (as of April 2026) were initially downloaded, totaling 252,456 entries. After filtering files containing at least one carbohydrate identifier in their “_chem_comp” data category structure, approximately 23,578 entries remained, representing 9.34% of the entire database. Nonetheless, this percentage is relatively low considering that a significant proportion of proteins in nature are glycosylated or interact with carbohydrates, emphasizing the under representation of carbohydrate-protein relationships in the database.^17,18^ The filtered entries are primarily derived from two experimental methods: X-ray crystallography (77.26%) and electron microscopy (22.14%) (Supplementary Figure 2). The main method of obtaining carbohydrate information in the PDB, X-ray crystallography, is limited by its reliance on the formation of regular crystals, which faces difficulties with flexible molecules such as carbohydrates.^19^ As a consequence, entries containing this family of biomolecules remain predominantly concentrated in medium to low resolutions (Table 2).

All glycan chains were then separated, and each individual carbohydrate monomer represented in the file structure was also isolated. This approach ensures that if an entry contained simultaneously a disaccharide and a tetrasaccharide as two independent chains, the data would be separated, and each of their monomers would be individually represented, resulting in six monosaccharide files. Additionally, all standalone carbohydrates were included. In cases where different conformations of a set of atoms were present, they were treated as distinct residues in a process that increased the amount of data accessible for analysis while maintaining the conformational information available in the database. Following this approach, a total of 319,961 monosaccharide files were generated, 313,919 hexopyranoses (98,1%) and 6,042 pentafuranoses (1,9%). Among the hexopyranose structures, 141,241 were obtained via X-ray diffraction and 166,538 via electron microscopy, together accounting for 98% of the dataset. X-ray diffraction entries are concentrated in high resolutions (2.5 Å or higher), while electron microscopy entries generally have medium to low resolutions (2.5 Å or lower) (Table 2). Within these data we observe that the PDB oligo/polysaccharide data remains highly concentrated on small chain sizes (Table 1) and residues types (Supplementary Table 1), as we previously observed,^11^ independently of recent advances on high throughput crystallography and CryoEM. Isolated monosaccharides represent more than half of all the PDB glycan structures (58,5%), followed by disaccharides (24,5%) and trisaccharides (8%). Only 9% of the sugar chains in the database are formed from four or more residues. Among these, approximately 51.1% are related to branched structures, while the remaining are linear saccharides. Notably, more than half of the carbohydrate entries contain at least one occurrence of N-acetyl-D-glucopyranosamine (D-GlcpNAc NDG/NAG). The second most frequent monosaccharide is D-mannopyranose (D-Manp MAN/BMA), followed by D-glucopyranose (D-Glcp GLC/BGC), reflecting more the focus of research and scientific interest than their occurrence in living organisms.

**Table 1:** Length of Glycan Chains in the PDB. Data for branched or linear saccharides were calulatd only for saccharides with 4 or more residues.

| Size | Quantity | Percent (%) |
| --- | --- | --- |
| Monosaccharide | 105730 | 59.88 |
| Disaccharide | 40960 | 23.2 |
| Trisaccharide | 14266 | 8.08 |
| Tetrasaccharide | 5049 | 2.86 |
| Pentasaccharide | 4570 | 2.59 |
| Hexasaccharide | 2437 | 1.38 |
| Heptasaccharide | 1530 | 0.87 |
| Eight or more | 2021 | 1.14 |
| **Total** | 176563 | - |
| **Linear** | 7029 (45.04%) |  |
| **Branched** | 8578 (54.96%) |  |

**Table 2:** Distribution of Chair and Non-Chair conformations by resolution in X-Ray Crystallography and Electron Microscopy^1^.

| Resolution (Å) | X-Ray Crystallography |  | Electron Microscopy |  |
| --- | --- | --- | --- | --- |
|  | Chair | Non-Chair | Chair | Non-Chair |
| $R \leq 1.0$ | 478 (96.4%) | 18 (3.6%) | 0 | 0 |
| $1.0 < R \leq 1.5$ | 6795 (91.6%) | 625 (8.4%) | 0 | 0 |
| $1.5 < R \leq 2.0$ | 34037 (90.3%) | 3638 (9.7%) | 705 (91.4%) | 66 (8.6%) |
| $2.0 < R \leq 2.5$ | 34724 (86.9%) | 5239 (13.1%) | 9868 (92.7%) | 774 (7.3%) |
| $2.5 < R \leq 3.5$ | 39419 (84.4%) | 7316 (15.6%) | 91524 (94.1%) | 5736 (5.9%) |
| $R > 3.5$ | 7711 (86.1%) | 1241 (13.9%) | 54630 (94.4%) | 3227 (5.6%) |

<sup>1</sup> The chair conformation was defined by cutoffs for ${}^4C_1$ ( $\theta \leq 26^\circ$ ) and ${}^1C_4$ ( $\theta \geq 154^\circ$ ).<sup>23</sup>

### Puckering

In solution, monosaccharides primarily adopt cyclic forms that can appear in many spatial conformations. Hexop0yranoses predominantly exhibit two main minima (^4^C_1_ and ^1^C_4_), corresponding to chair conformations, along with several transitional states or local minima: Boat (B), Skew-Boat (S), Envelope (E), and Half-Chair (H). Chairs are generally the preferred conformations for saturated rings, as they allow the optimal 60^*°*^ angle between the substituents. Due to the need of specular movements during the cyclization of D-and L-sugars, and save for few exceptions, D-sugars typically find their main minima in a ^4^C_1_ conformation, whereas L-sugars often adopt a ^1^C_4_ conformation.^17,20^ While the energy barriers separating each conformation are significant, non-chair conformations have been reported as important intermediate states in various biological processes, such as hydrolysis, glycoside transfer from an activated nucleotide-sugar, isomerization, and epimerization.^20^ Thus, to acquire a clear picture of the extent in which PDB has captured carbohydrate puckering in biological systems, we employed Cremer-Pople puckering coordinate measurements for all hexopyranoses deposited in the database. As shown in Table 2, both X-ray diffraction and cryoelectron microscopy data reveal a direct relation between resolution and the observation of non-chair conformations for hexopyranoses. However, in Cryo-EM data, the observation of such high energy states is up to ten times lower in magnitude when compared to X-ray crystallography, evidencing the role of each method and experimental conditions in the observation of monosaccharide distortions. Reinforcing this assumption, analysis of the real-space correlation coefficient demonstrated that the percentage of high-energy conformations also increases in structures that exhibit poor similarity between electron-density and experimental data (Figure 1). To further assess these correlations, a subset of high-resolution pyranose structures was created. To ensure the quality of the structures, we selected those with a resolution of 2.0 Å or better and an Occupancy Weighted Average B factor (OWAB) of 60 Å or less.^11,21,22^

**Figure 1:**
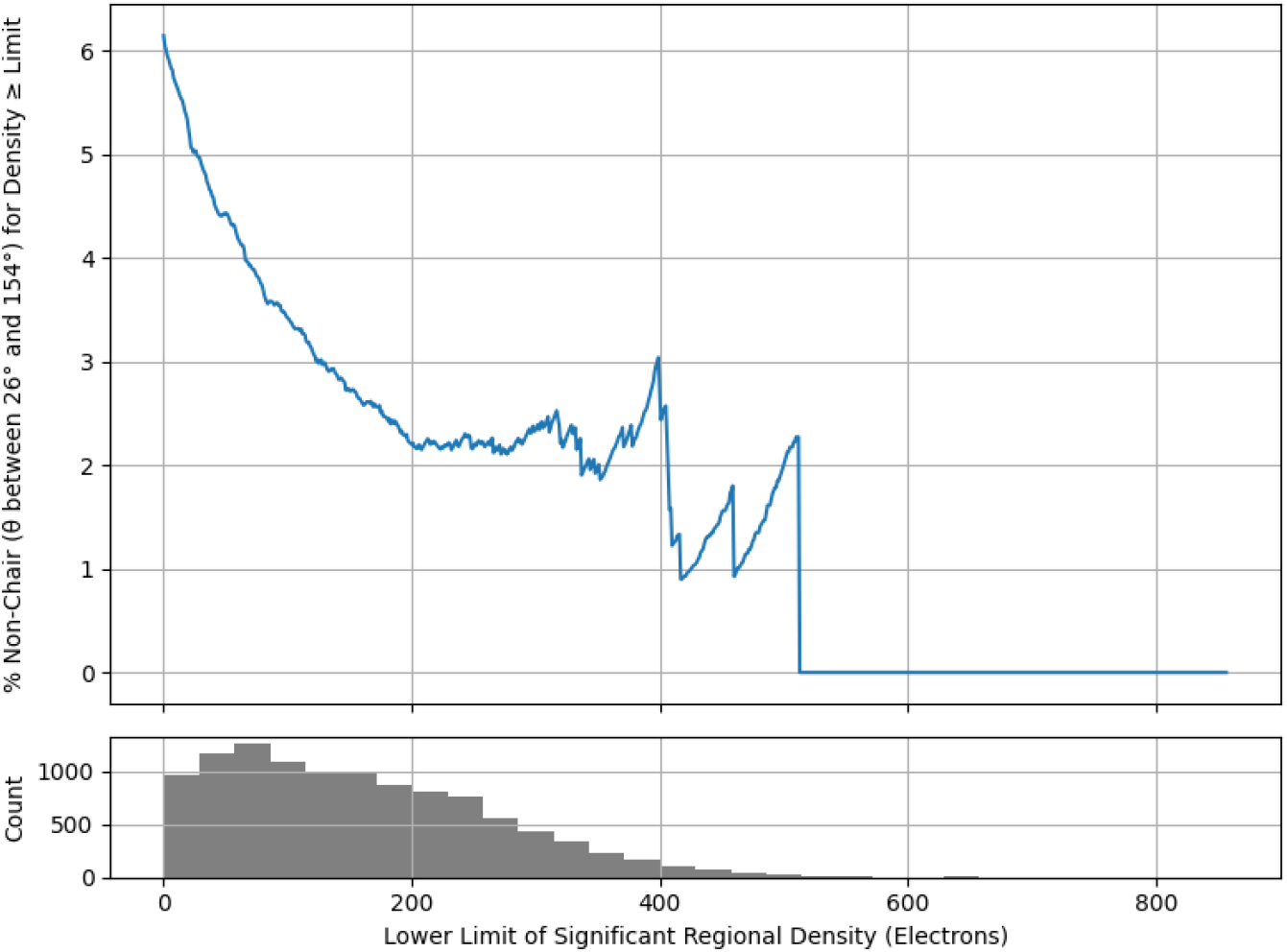
Cumulative percentage of non-chair conformations as a function of the structural similarity to the electron density data, demonstrating that hexopyranoses distortion is highly correlated with the experimental data quality.

On average, the PDB shows 6.0% of hexopyranose residues in non-chair conformations, ranging from 4% to 7% among the five main residues. So there is no correlation between the level of distortion and the monosaccharide identity (Figure 2). Also, such conformations are mostly high energy states, not located on local minima of the residues (the main exception here seems to be mannose). These observations suggest that the effects interfering with residue distortions are, on average, similar among all residues, which points to a more likely experimental effect than a specific protein-induced effect. However, while this experimentally average behavior is able to explain the residues distortions associated with proteins without catalytic activity, the specific influence of enzymes in complex with substrates are likely to be hidden in the average. So, to circumvent this limitation and better explore how non-chair conformations are distributed in biomolecular systems, we intersected the PDB carbohydrate information with the Carbohydrate-Active Enzymes (CAZy, Table 3).

**Figure 2:**
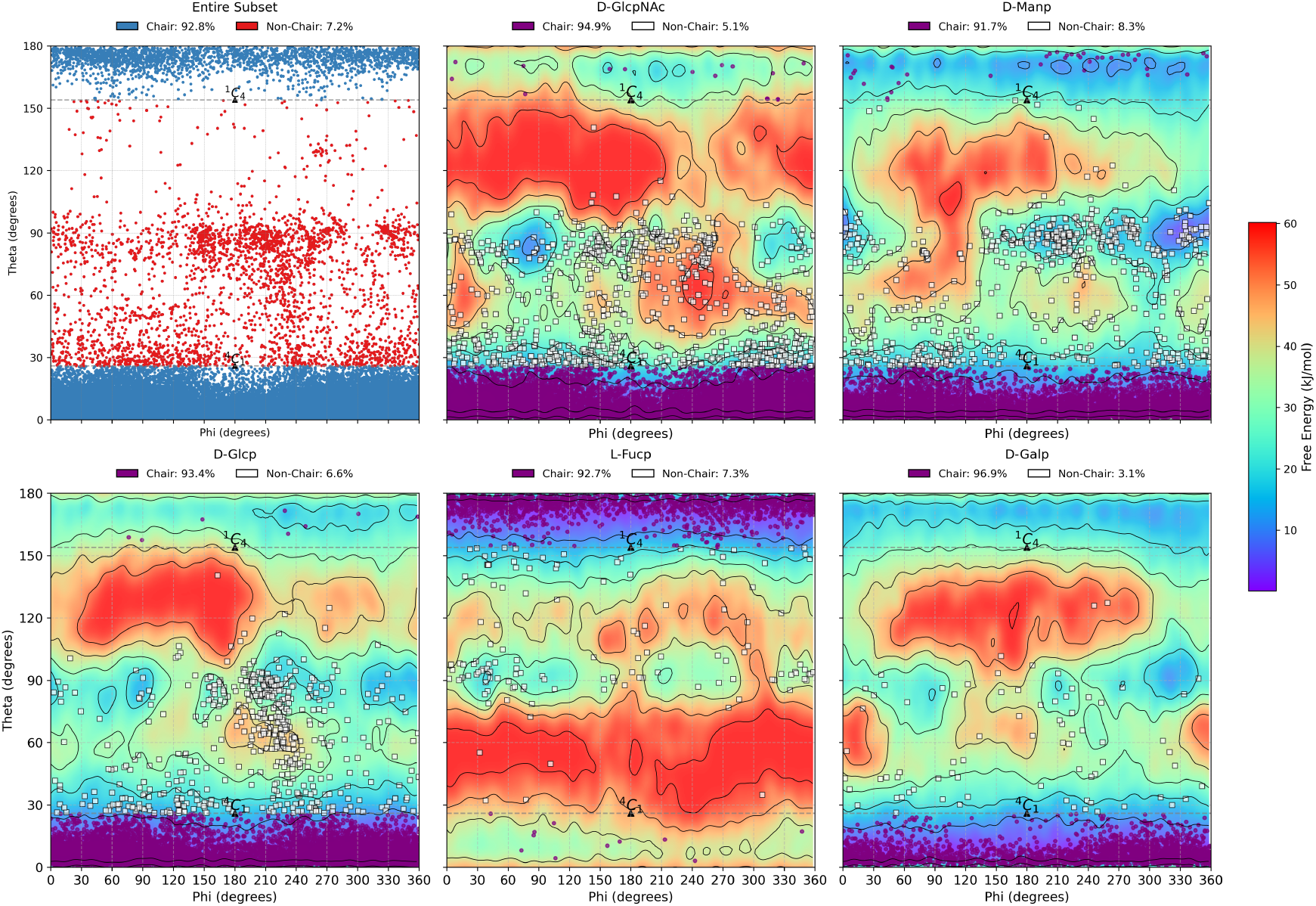
Conformational free energy landscapes from QM/MM metadynamics simulations and puckering coordinate values for the pyranose subset, highlighting the five most abundant monosaccharides found in the PDB. The chair conformation was defined by cutoffs for ^4^C_1_ (*θ* ≤ 26°) and ^1^C_4_ (*θ* ≥154°).^23^ The *α* and *β* anomers are considered together. QM/MM simulations were performed using the PBE functional with a DZVP basis set. For clarity, chair conformations are shown as purple circles, whereas non-chair conformations are shown as white squares.

**Table 3:** Distribution of chair and non-chair conformations among carbohydrate-related enzyme families.

| Family | CHAIR | NON-CHAIR | Percent (%) |
| --- | --- | --- | --- |
| Glycoside Hydrolases | 13,653 | 1296 | 8.67 |
| Carbohydrate-Binding Modules | 2,508 | 149 | 5.61 |
| Auxiliary Activities | 1,285 | 68 | 5.03 |
| Glycosyltransferases | 630 | 44 | 6.53 |
| Polysaccharide Lyases | 430 | 81 | 15.85 |
| Carbohydrate Esterases | 256 | 4 | 1.54 |

The organization of monosaccharides puckering following CAZy classification (Figure 3) maintains the magnitude of distortion within the PDB average, between 1% and 7%, among glycoside hydrolases (GHs), carbohydrate-binding Modules (CBMs), auxiliary activities enzymes (AAs), glycosyltransferases (GTs), carbohydrate esterases (CEs), and polysaccharide lyases (PLs). This range of values is probably influenced by the largely different number of representatives in each class (Table 3). Still, the enzyme class most represented in the PDB is the one with the largest number of distorted residues (GHs, ~ 7%). As a reference, the CBM group, which lacks catalytic activity, exhibits a smaller proportion of high-energy conformations, but still within the average behavior of the database. Additionally, the largely different number of entries in each family poses a challenge for the establishment of class specific properties.

**Figure 3:**
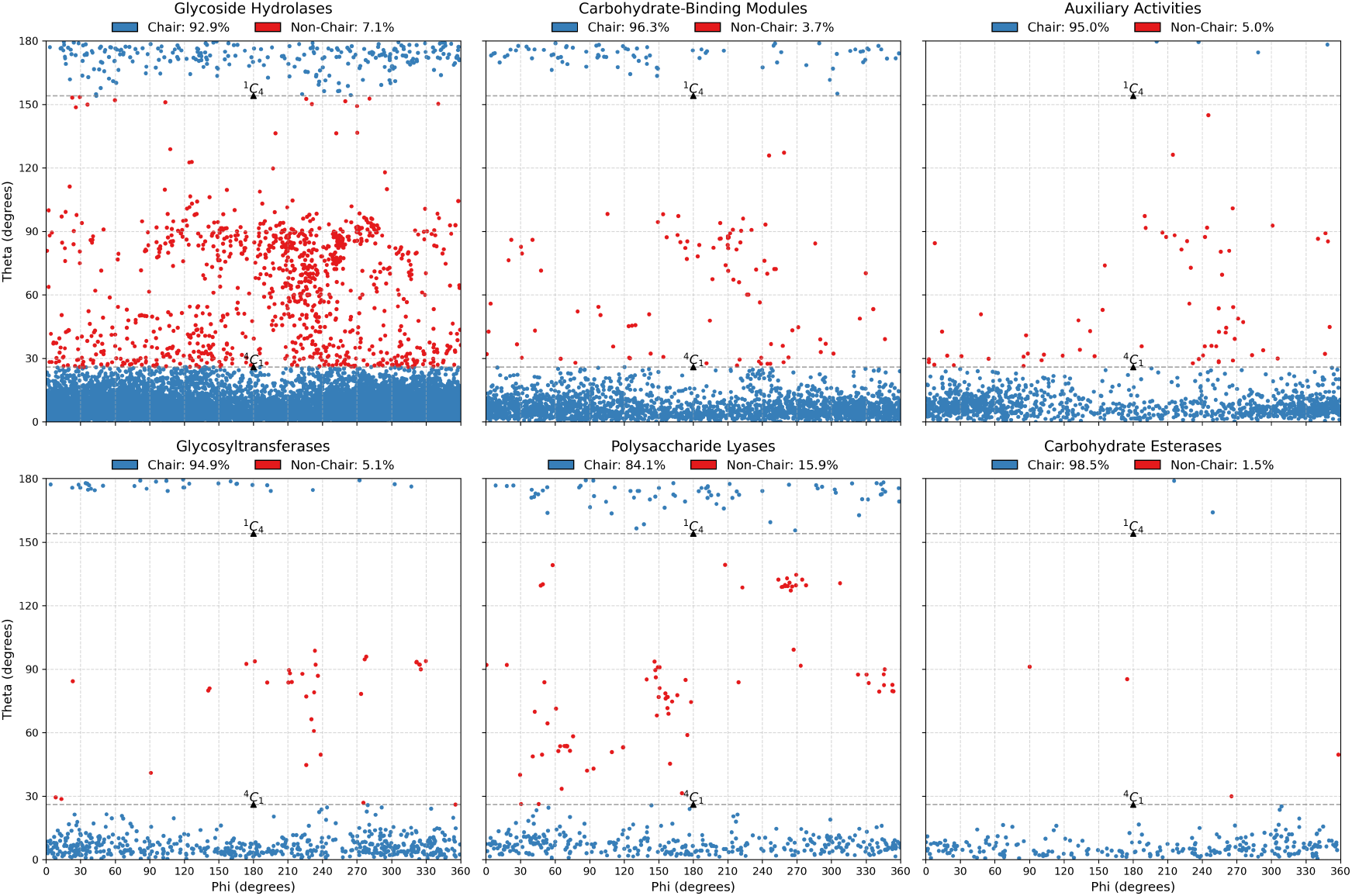
Puckering coordinate values for the pyranose subset, grouped by CAZyme class. The chair conformation was defined by cutoffs for^4^C_1_ (*θ* ≤ 26°) and^1^C_4_ (*θ* ≥154°).^23^ The *α* and *β* anomers are considered together.

In previous studies, the sugar ring conformations^1^,^4^B (*ϕ* = 240° and *θ* = 90°) and ^1^S_3_ (*ϕ* = 210° and *θ* = 88°) were identified as local minima for N-acetyl-D-glucosamine, with these states suggested to be induced by Glycoside Hydrolases (GH). In the subset, a total of 37 and 53 structures adopted the ^1^,^4^B and ^1^S_3_ states, respectively, with 62 ( ≈ 69%) of these originating from GH entries. While this may suggest a correlation, it is important to note that these two states account for only 11% of all non-chair conformations observed in the subset for N-acetyl-D-glucosamine.^11,24–26^

Similarly, for D-glucopyranose, studies have described an itinerary involving ^1^S_3_,^4^H_3_ (*ϕ* = 240° and *θ* = 90°) and ^4^C_1_ as a significant pathway used by GHs during catalysis.^27,28^ Of all D-glucopyranose structures, 36 adopted the ^4^H_3_ conformation and 86 adopted the ^1^S_3_ conformation, with 61 (50%) of these belonging to GH entries. Together, these two conformations represent 12.6% of all non-chair conformations observed for D-glucopyranose in the subset. For other monosaccharides analyzed in Figure 2, non-chair conformations Swere also identified and correlated with enzyme pathways. For example,^1^S_3_ and^1^S_5_ were observed for D-mannopyranose (associated with mannosidases),^29,30^ ^1^S_3_ was observed for D-galactopyranose (associated with galactosidases),^31,32^ and ^3^S_1_ was observed for L-fucopyranose (associated with fucosidases).^11,33,34^ Taken together, these results suggest that while most of the carbohydrate rings’ deformations found in the PDB are likely experimental artifacts, GHs in particular seem to be able to interfere with the hexopyranoses conformational space, inducing non-chair conformations. In order to explore the molecular basis for these observations, we proceeded to measure the free energy landscapes (FELs) of different residues complexed with proteins, to check the ability of different proteins environments to interfere in carbohydrates conformational landscape.

### Exploring the energetics of carbohydrates

To access the ability of proteins to interfere in carbohydrate residues conformational space, possibly stabilizing high-energy, non-chair conformations, we performed a series of metadynamics calculations on complexes of carbohydrates and various CAZYmes. Figure 4 shows the FELs of bound monosaccharides in complex with their respective proteins, grouped by CAZyme classes:

- D-Xylp in complex with CBM X-2 L110F from *Rhodothermus marinus* (5DPN; 1.60 Å);
- D-Glc in complex with AMP-activated protein kinase CBM from *Rattus norvegicus* (1Z0N; 1.49 Å);
- L-Arap in complex with arabinoxylanase from *Acetivibrio thermocellus* (5LA2; 1.65 Å);
- D-GlcNAc in complex with EndoBT-3987 from *Bacteroides thetaiotaomicron VPI-5482* (6TCV; 1.31 Å);
- D-Glc in complex with GlgE from *Mycobacterium thermoresistibile* (5CGM; 1.95 Å);
- D-Glc in complex with Lichenase F32EG5 from *Caldicellulosiruptor sp. F32* (5H4R; 1.70 Å);
- D-Manp in complex with MPT-2 from *Leishmania mexicana* (6Q4Z; 1.55 Å);
- D-Glc in complex with Spinosyn Rhamnosyltransferase from *Saccharopolyspora spinosa* (3UYL; 1.85 Å);
- D-Glc in complex with PHS2 from *Arabidopsis thaliana* (4BQI; 1.90 Å).

**Figure 4:**
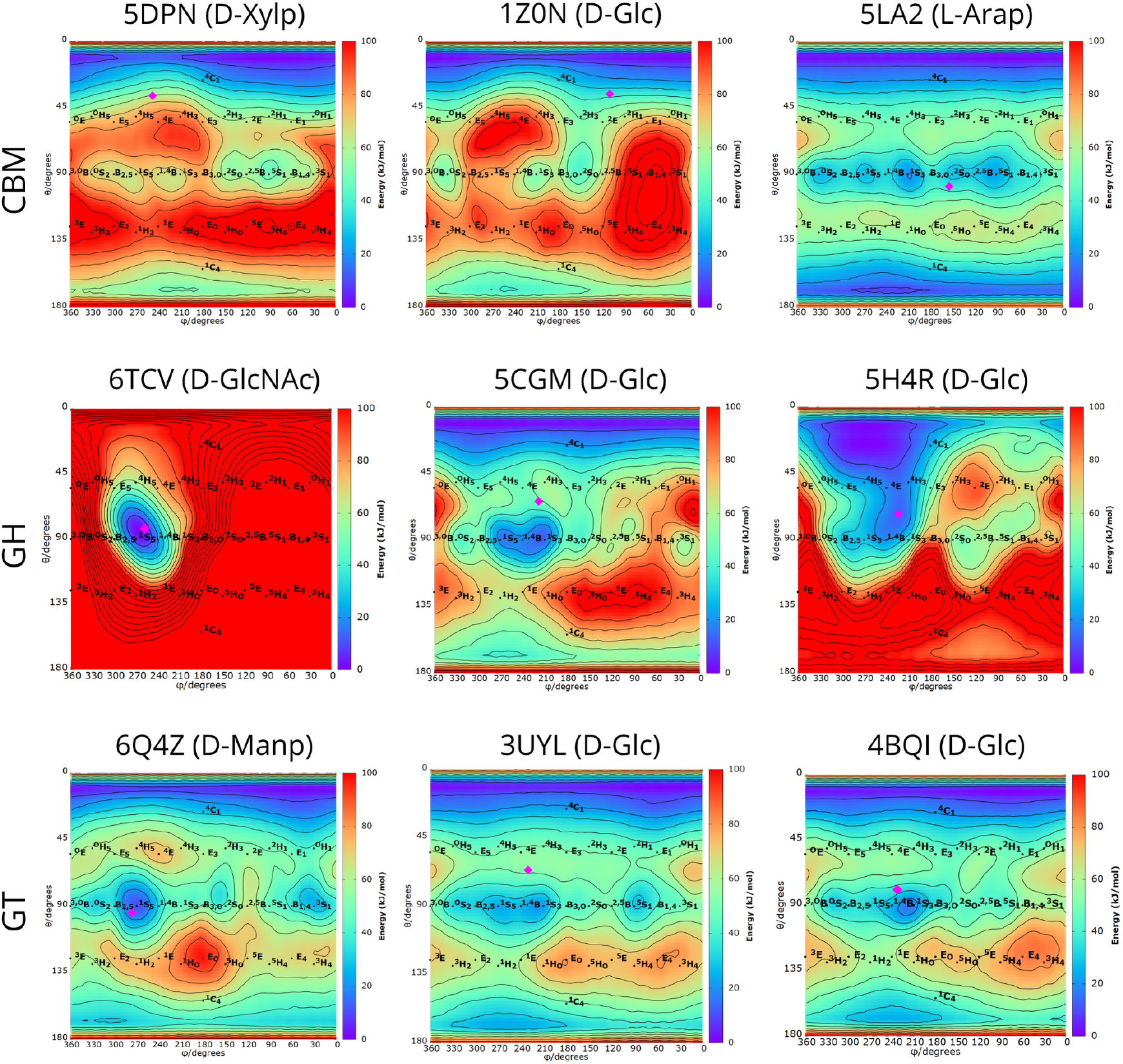
Puckering free energy landscapes (FEL) of monosaccharides in sugar-protein complexes obtained from metadynamics simulations. The puckering coordinates of each isolated sugar are highlighted in magenta. Panels are grouped by CAZyme class: carbohydrate binding modules (top row), glycoside hydrolases (middle row) and glycosyltransferases (bottom row). PDB entry ID and sugar names are presented above each panel. The metadynamics simulations were performed with protein side chains left unrestrained, using the CHARMM3m force field and the *ϕ* and *θ* torsion angles as colective variables. The chair conformation was defined by cutoffs for ^4^C_1_ (*θ* ≤ 26°) and ^1^C_4_ (*θ* ≥ 154°).^23^

We observed that, in most cases, the protein scaffold is not able to stabilize non-chair conformations upon complexation (Figure 4), with the exception of GHs. These enzymes showed distinct behaviors during simulations, from the absence of influence (PDB ID 5CGM) to medium (PDB ID 5H4R) and large (PDB ID 6TCV) stabilization of non-chair states. Particularly in the case of PDB ID 5H4R, we observe that, despite the stabilization of the equatorial regions of the map, the global minimum remains on the expected chair conformation. The only situation in which we observed the ability of a protein to unequivocally define the crystallographic non-chair state as the global minimum is in PDB ID 6TCV. It’s important to emphasize that these results were obtained in high resolution structures (<2Å), so different aspects from the experimental conditions, such as non-native temperatures, concentrations, presence of non-native compounds, crystal packing, cryogenic conditions or methodological aspects such as force fields and structural refinement steps could be operating in most situations with distorted residues. In other words, even high quality structures, with well defined electron densities, may still offer carbohydrate conformations with no biological significance.

In a previous work, our group analyzed the structure of the CBM21 from *Rhizopus oryzae* glucoamylase (RoGA) (PDB entry code 4BFN, 1.34 Åresolution) that contains a non-chair carbohydrate conformation (^1^S_3_ *α*-D-Glc). Through a similar methodological approach, through metadynamics to sample carbohydrates conformational space with and without the protein influence, we observed that the experimental distorted state is likely an artifact and not biologically relevant for CBM21 complexation or recognition of isomalto oligosaccharides,^35^ highlighting the challenge of assuming potential biological relevance for generally unusual conformations observed in the PDB. This challenge gains importance considering the relevance of carbohydrate binding proteins, such as CBMs, in multiple industrial sectors, including the optimization of food industry processes,^36^ enhanced biomass degradation^37–39^ and cellulose fiber processing.^40^

In the search of a better understanding of protein-carbohydrate interactions, able to improve the design of both bioactive compounds and the development of predictive models, the current work presented a comprehensive overview analysis of carbohydrate information within the PDB, implementing standardized carbohydrate annotation and multiple structure quality filters. This information was classified, quantified, and presented across all carbohydrate-containing entries in the database. By incorporating Cremer-Pople puckering coordinates, we characterized all separated hexopyranose conformations. Our study reveals that 3D structural carbohydrate data remains highly concentrated in small glycan chains and a limited number of sugar residues, with more than 97% of the data obtained through X-ray diffraction or electron microscopy (with a substantial growth of the latter in the recent years). Data indicates a correlation between unusual conformational states and structural quality, highlighting the need for caution in assuming biological relevance for the non-chair conformations observed in the PDB. In this regard, metadynamics simulations have proven to be an effective tool to explore the role of the protein binding site in the possible modification of carbohydrate residues conformational landscape. Finally, the presented combination of structural analysis and simulations reinforce the hypothesis that carbohydrate distortions on PDB are mostly derived from the experimental conditions, without biological implications, while a handful of enzymes, particularly GHs, seem to be able to stabilize particular non-chair conformations.

## Methods

### Data Retrieval

All PDB entries were downloaded on April 28, 2026, in mmCIF format and filtered to identify carbohydrates with the assistance of the Bio.PDB.MMCIF2Dict module of the BioPython library.^41^ Entries containing one or more carbohydrates in their chemical components list (_chem_comp.id) were selected, considering the carbohydrate IDs provided by the wwPDB carbohydrate remediation (CCD carbohydrate list).

### Sugar separation

For each carbohydrate-containing entry, all sugar components listed in the CIF data category “atom site” were isolated and treated as individual structure files for puckering measurements. Additionally, information such as their IUPAC carbohydrate symbols and entity types (branched or non-polymer) was gathered for analysis. Each structure file was named using a standardized format comprising the PDB Entry ID, Sugar Component ID, and the first atom number from the “atom site” category, separated by underscores. For instance, the N-acetyl-D-glucopyranosamine from the PDB entry ID 1UHG, with its first atom number being 11983, was named “1uhg_NAG_11983”. Finally, all information for entities classified as ‘branched’ in the CIF entity data category was extracted to identify glycan chains.

### Puckering

The separation process resulted in PDB files containing individual monosaccharides, including monomers extracted from oligosaccharides. The PUCKERING function of PLUMED Driver was used to calculate the pseudorotational coordinates of the rings in each generated PDB file.^42^ For each pyranose, the atom numbers forming a ring were identified in the program’s required order (O5, C1, C2, C3, C4, and C5), and the *θ* and *ϕ* pseudorotation coordinates were computed. To distinguish Chair from Non-Chair conformations, *θ* angles of 26°(^4^*C*_1_ chair) and 154°(^1^*C*_4_ chair) were used as cutoffs.^43^ Non-Chair conformations were characterized based on their proximity to canonical conformation coordinates.^23^

### Molecular Dynamics

We selected the structures of the five most prevalent sugars in the PDB, along with ten CAZy-ranked complexes, chosen based on their high electron density values, that exhibited distorted hexopyranose rings. This selection aimed to provide a comprehensive view of how sugar ring puckering behaves in aqueous solution compared to their conformations in crystalline environments. The selected structures were processed using CHARMM-GUI,^44^ and system topologies were generated with the CHARMM36m force field for GROMACS.^45,46^ Molecular dynamics (MD) simulations were conduted using GROMACS 2024.^47^ The systems equilibration followed a protocol previously described by our group,^35,48^ starting with energy minimization step using the steepest descent algorithm, followed by a 1 ns NVT equilibration with position restraints of 5,000 kJ mol^−1^ applied to solute heavy atoms. This was followed by six NPT equilibration steps. The first NPT step maintained the same restraints as the NVT phase, while in each subsequent step the restraint force was reduced by 1,000 kJ mol^−1^. In the final (sixth) step, restraints of 1,000 kJ mol^−1^ were applied only to the solute backbone, allowing side chains to move freely. During simulations the LINCS algorithm was used to constrain bonds involving hydrogen atoms,^49^ allowing a simulation timestep of 2 fs. During equilibration, the C-rescale barostat (*τ* = 2 ps) was coupled with the V-rescale thermostat (*τ* = 0.1 ps) to maintain constant pressure and temperature.^50,51^ For production simulations, the Parrinello–Rahman barostat (*τ* = 2 ps) was used instead.^52^ Nonbonded interactions were calculated using a 1.2 nm cutoff, and long-range electrostatics were treated with the Particle Mesh Ewald (PME) method.^53,54^

### Quantum Mechanics/Molecular Mechanics

We combined Quantum Mechanics/Molecular Mechanics (QM/MM) and metadynamics methods to determine the puckering free energy profiles of the most prevalent monosaccharides in the PDB as a reference for their expected behavior in solution: D-GlcNAc, D-Manp, D-Glcp, L-Fucp, and D-Galp. For each system, a snapshot taken at the end of the equilibration phase was used as the starting structure for the QM/MM simulations. Simulations were performed using the interface between the CP2K^55^ and GROMACS^47^ codes. All QM calculations were conducted within the density functional theory (DFT) framework, using the Perdew–Burke–Ernzerhof (PBE) generalized gradient approximation.^56^ The Goedecker–Teter–Hutter (GTH) dual-space pseudopotentials^57^ were used in conjunction with the Gaussian and Plane Wave (GPW) scheme,^58^ as implemented in the CP2K/Quickstep module,^59^ enabling accurate and efficient DFT calculations for large systems. For each system, a QM region encompassing all atoms of the monosaccharide was defined. A DZVP-MOLOPT-GTH basis set was used, with a plane-wave cutoff of 450 Ry. The interactions between the QM and MM subsystems were treated using the Gaussian Expansion of the Electrostatic Potential (GEEP) method developed by Laino et al.^60^ The simulations were run with a timestep of 0.12 fs for a total of 240 ps. The size of the QM box was defined as 10 × 10 × 10 Å. The self-consistent field (SCF) convergence criterion was set to 1.0×10^−6^.

### Metadynamics

To investigate how sugars pucker in solution and in crystallographic complexes, we performed well-tempered metadynamics simulations using PLUMED 2.9,^61,62^ allowing us to determine the free energy landscape (FEL) of hexopyranose rings along the Cremer–Pople puckering coordinates *ϕ* and *θ*.^63^ Metadynamics simulations were conducted using *ϕ* and *θ* as collective variables (CVs), with Gaussian heights of 1 kJ mol^−1^ for MM simulations and 2.5 kJ mol^−1^ for QM/MM simulations. The Gaussian width was set to 0.1 rad, a bias factor of 10 was used, and a new Gaussian-like potential was deposited every 200 MD steps. During simulations of protein–glycan complexes, a harmonic positional restraint of 1,000 kJ mol−1 was applied to the protein backbone and glycan residues not involved in the metadynamics bias. This approach ensured that only the FEL of the distorted sugar residue observed in the crystallographic complex was sampled. By preventing conformational changes in the rest of the glycan, it avoided sampling geometries inconsistent with the crystallographic structure and thereby prevented distortion of the FEL.

## Acknowledgments

The authors would like to thank CAPES, CNPq, FAPERGS, and the National Laboratory for Scientific Computing (LNCC/MCTI, Brazil) for the use of the Santos Dumont supercomputer.

## Conflict of Interest

The authors declare no competing financial interests

## Notes

### Competing Interest Statement

The authors have declared no competing interest.

### Funding

- Coordenação de Aperfeicoamento de Pessoal de Nível Superior
- Conselho Nacional de Desenvolvimento Científico e Tecnológico
- Fundação de Amparo à Pesquisa do Estado do Rio Grande do Sul
- Laboratório Nacional de Computação Científica

