## Supplemental material for "Are carbohydrate ring distortions determined by biology or crystallography? The impact of experimental conditions on structural data"

### Supplementary Information

Table 1: Most commonly found monosaccharide residues in PDB entries.

| Name | Entries |
|---|---|
| D-GlcpNAc | 15576 |
| D-Manp | 6487 |
| D-Glcp | 3432 |
| L-Fucp | 2514 |
| D-Galp | 1843 |
| D-Xylp | 465 |
| D-GalpNAc | 423 |
| D-Glcp6PO3 | 144 |
| L-Rhap | 100 |
| D-GlcpNSO36SO3 | 90 |
| Others | 2294 |

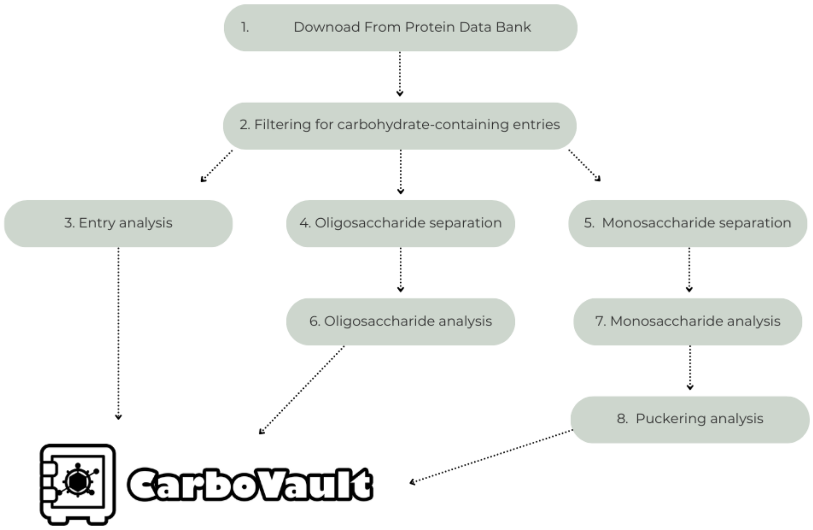


Figure 1: Workflow used to collect and analyze carbohydrate data from the PDB. The code is available on request from the authors.

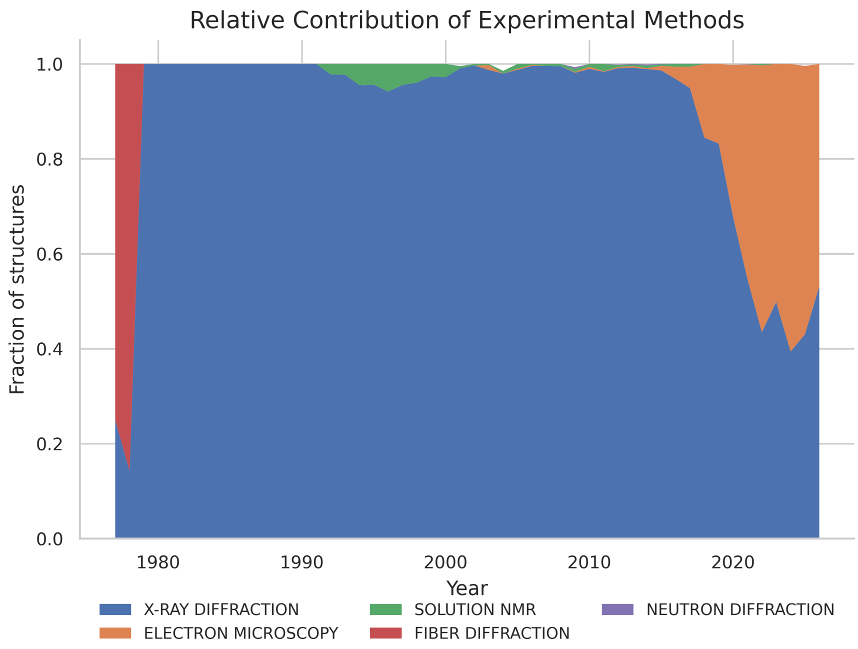


Figure 2: Fractional contribution of experimental methods over time for Protein Data Bank entries containing saccharides, showing changes in method prevalence across deposition years.
